# Fusion pore expansion and contraction during catecholamine release from endocrine cells

**DOI:** 10.1101/2020.03.02.972745

**Authors:** M. B. Jackson, Y.-T. Hsiao, C.-W. Chang

**Affiliations:** U. of Wisconsin Med. Sch.; University of Wisconsin; Gladstone Inst UCSF

## Abstract

Amperometry recording reveals the exocytosis of catecholamine from individual vesicles as a sequential process, typically beginning slowly with a pre-spike foot, accelerating sharply to initiate a spike, reaching a peak, and then decaying. This complex sequence reflects the interplay between diffusion, flux through a fusion pore, and possibly dissociation from a vesicle’s dense-core. In an effort to evaluate the impacts of these factors, a model was developed that combines diffusion with flux through a static pore. This model recapitulated the rapid phases of a spike, but generated relations between spike shape parameters that differed from experimental results. To explore the possibility of fusion pore dynamics, a transformation of amperometry current was introduced that yields fusion pore permeability divided by vesicle volume (*g/V*). Applying this transform to individual fusion events yielded a highly characteristic time course. *g/V* initially tracks the pre-spike foot and the start of the spike, increasing ∼15-fold to the peak. However, after the spike peaks, *g/V* unexpectedly declines and settles to a constant value that indicates the presence of a stable post-spike pore. *g/V* of the post-spike pore varies greatly between events, and has an average that is ∼3.5-fold below the peak value and ∼4.5-fold above the pre-spike value. The post-spike pore persists and *g/V* remains flat for tens of milliseconds, as long as catecholamine flux can be detected. Applying the *g/V* transform to rare events with two peaks revealed a stepwise increase in *g/V* during the second peak. The *g/V* transform offers an interpretation of amperometric current in terms of fusion pore dynamics and provides a new framework for analyzing the actions of proteins that alter spike shape. The stable post-spike pore conforms with predictions from lipid bilayer elasticity, and offers an explanation for previous reports of prolonged hormone retention within fusing vesicles.

**STATEMENT OF SIGNIFICANCE:** Amperometry recordings of catecholamine release from single vesicles reveal a complex waveform with distinct phases. The role of the fusion pore in this waveform is poorly understood. A model based on a static fusion pore fails to recapitulate important aspects of the waveform. A new transform of amperometric current introduced here renders fusion pore permeability in real time. This transform reveals rich dynamic behavior of the fusion pore as catecholamine leaves a vesicle. This analysis shows that fusion pore permeability rapidly increases and then decreases before settling into a stable post-spike configuration.

## INTRODUCTION

Amperometry recording from catecholamine-secreting endocrine cells reveals the exocytosis of single vesicles (1). At a course-grained level a single-vesicle appears to release its catecholamine content in an instantaneous burst. Closer inspection reveals a succession of stages that are thought to reflect a number of different factors, including flux through a fusion pore, diffusion from the cell surface to the electrode surface, and dissociation from the protein matrix of an endocrine vesicle’s dense-core. The earliest phase of release that is visible in an amperometry recording is the pre-spike foot (2, 3), which has been studied to gain insight into the nature of the initial fusion pore (4, 5). This initial stage is followed by the onset of a spike as the fusion pore expands. Once this expansion has started it becomes very difficult to tease apart the various contributions, and the role of the fusion pore becomes difficult to define. The rise of the spike is so rapid that it appears to be diffusion limited, and the influence of the fusion pore may not be detectible. The decay of the spike is too slow to be diffusion limited, and dissociation of catecholamine from the proteins of the vesicle matrix has been invoked to account for this feature (3, 6-8). However, a role for expanding fusion pores is implied by a large number of studies reporting that many fusion proteins influence the spike waveform (9-19). Most of these proteins reside in the cytoplasm, out of reach of the matrix within a vesicle. To alter the shape of a spike these proteins must interact with a vesicle from the outside, presumably with membrane at or near the fusion pore. These results thus support the idea that the fusion pore has a significant role in shaping an amperometric spike.

Fusion pores can also limit the release of peptide hormones tagged with fluorescent proteins (20-22). The prolonged retention of these large molecules within a vesicle after the onset of fusion suggests that fusion pores remain narrow for seconds. Furthermore, the selective release of small molecules alone or small molecules together with large molecules indicate that fusion pore expansion is subject to biological control (23-25).

Although amperometry recording has illuminated many aspects of exocytosis in remarkable detail, using this technique to study the progress of fusion beyond the pre-spike foot has proven challenging. Changes in spike shape have often been interpreted qualitatively in terms of alterations in expanding fusion pores, but the quantitative relationship between fusion pores and amperometric spikes is not well understood. The present study explores this relationship. A model incorporating diffusion and flux through a static pore was developed and its predictions tested against experiment. A static fusion pore model could fit the fast components of a spike, but failed to recapitulate observed correlations between shape indices. Turning to a dynamic representation of the fusion pore lead to a new transformation of amperometry data that yields fusion pore permeability divided by vesicle volume. This approach suggests that an initial rapid component of spike decay represents a change in the fusion pore. A slow component of spike decay represents flux through a stable post-spike fusion pore. The dynamic fusion pore offers a new perspective on amperometric spikes and suggests that the post-spike pore serves as a locus for control by proteins.

## EXPERIMENTAL METHODS

This study employed standard methods of amperometry recording from chromaffin cells, as reported previously from this laboratory (26). Adrenal glands were dissected from newborn wild type mice, dissociated, and cultured in DMEM containing penicillin/streptomycin and insulin-transferrin-selenium-X on poly-d-lysine coated cover slips following the procedure of Sorensen et al (27). Amperometry recordings were made from cells 1-3 days *in vitro* at room temperature (∼22 °C) with a bathing solution consisting of (in mM) 150 NaCl, 4.2 KCl, 1 NaH_2_PO_4_, 0.7 MgCl_2_, 2 CaCl_2_, 10 HEPES (pH 7.4). A carbon fiber electrode was positioned to gently touch a cell, as this has been shown to minimize the time for diffusion from the release site to the electrode surface (6). Signals were amplified with a VA-10 amplifier (ALA Scientific Instruments). Exocytosis was triggered by pressure application of a depolarizing solution (bathing solution with 105 mM KCl and reduced NaCl) for 6 seconds. Records were acquired with an intel-based computer running PCLAMP software. Amperometric current was low-pass filtered at 2.5 kHz, and digitized at either 4 or 10 kHz.

Data was analyzed with the computer program Origin and with in-house software written in C++. Analysis and modeling were performed with the computer program Mathcad.

## RESULTS

### Spike Properties

Amperometric current reveals many spikes occurring in a 23 second recording episode, during and after the application of the depolarization solution (Fig. 1A). Three typical events in this long recording, marked by a small horizontal bracket above the 8 sec time point, are displayed on an expanded time scale in Figs. 1B1, 1B2, and 1B3. In each trace an exponential was fitted to the top 2/3 of the decay, revealing a slower component. This biphasic decay was evident in ∼95% of the events. The slow component had a relatively small and variable amplitude but it was generally clear. This slow component is comparable to the “post-spike foot” described by Mellander et al (28). However, that study highlighted events with roughly stepwise ends. In the present study the post-spike feet generally decayed exponentially to baseline. The decay of the slow component will be explored further below, and the modeling efforts will attempt to illuminate each phase of decay separately. At this point it is worth noting that the biphasic nature is qualitatively what one would expect for rapid release of a free fraction of catecholamine, followed by the slow release of a fraction bound to the vesicle matrix. From this perspective the rapid phase of decay should be limited by flux through a pore.

**Fig. 1.**
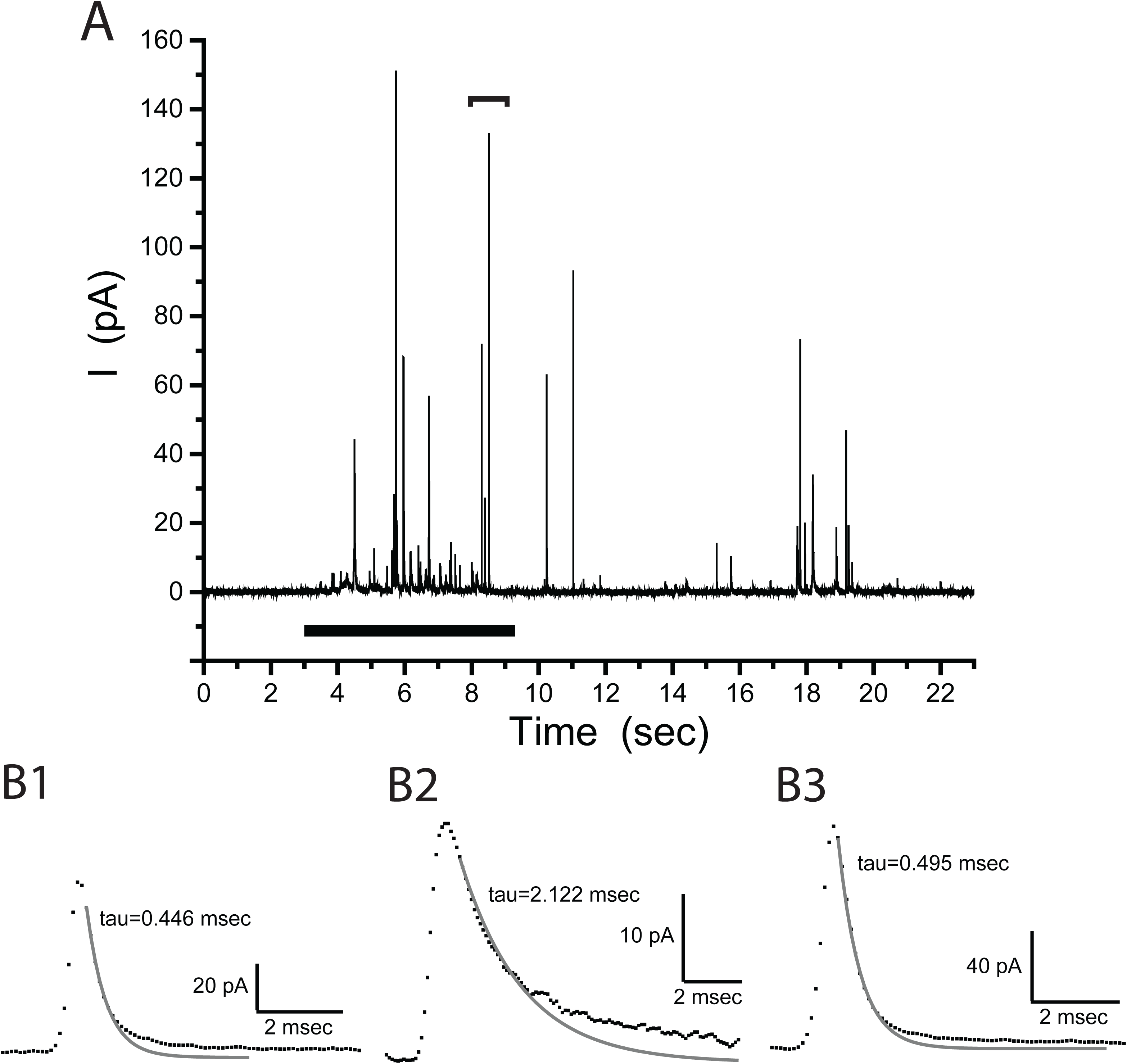
Amperometry recordings of single-vesicle release events in chromaffin cells. **A.** One long trace of current versus time illustrates that depolarization (thick horizontal bar from 3-9 seconds) elicits many events with variable amplitudes. The bracket above at ∼8 sec indicates three events displayed on an expanded scale in **B.** These three selected events occurred in sequence and illustrate variations in amplitude and kinetics. Single exponentials were fitted to the decaying phase within 2/3 of the peak. These fits are shown as gray curves. The later phase of current is clearly above the exponential curve fitted to the initial phase of decay. This highlights the two components of decay, the second of which is referred to as the post-spike foot (28).

For each event we determined the peak amplitude, 35-90% rise time, time constant of the rapid component of decay (from fits such as in Fig. 1B1, 1B2, and 1B3), and the number of molecules released, *N*_*0*_ (the integral of the amperometric current over an entire event gave the total charge, which was converted to molecule number by assuming two electrons per catecholamine molecule, and multiplying by Faraday’s constant). Measurements from 1682 events from 77 cells (digitized at 4 kHz) were used to construct distributions (Fig. 2). Their means and standard deviations are stated in the legend of this figure. The distributions in Fig. 2 illustrate the variations in all of the quantities, making the point that spike shape is quite variable. The absence of events below 20 pA in Fig. 2A reflects the use of an amplitude cutoff in the analysis in order to focus on events that are large enough to measure accurately. The rise times of < 1 msec are consistent with the close contact between the electrode and cell, which minimizes the diffusion time (6). It should be noted that kiss-and-run caused by fusion pore closure would prevent the release of the entire content, resulting in an apparent *N*_*0*_ value lower than the total number of molecules contained within a vesicle.

**Fig. 2:**
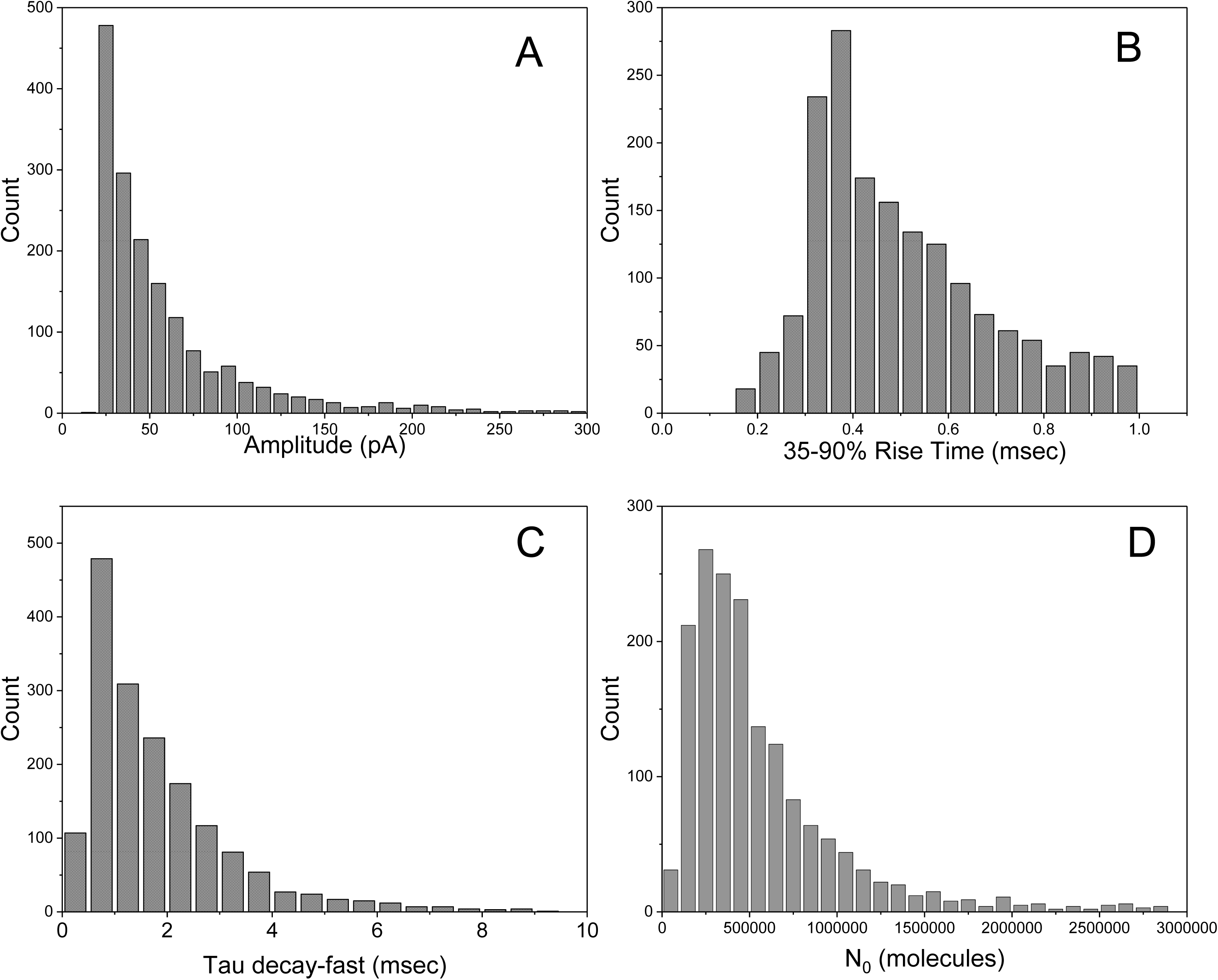
Distributions of **A.** peak amplitudes, **B.** 35-90% rise times, **C**. decay times of the fast component (Tau decay-fast; from fits as in Figs. 1B1-1B3), and **D.** number of molecules N_0_, computed from the total area of the event (charge converted to molecules with Faraday’s constant assuming two electrons per molecule of catecholamine). Means ± standard deviations are, peak amplitude 59.9 ± 50.7 pA, 35-90% rise time 0.50 ± 0.19 msec, Tau decay-fast 1.84 ± 1.50 msec, N_0_ 600,080 ± 611,222. Plots were based on 1682 events recorded from 77 cells.

### Static Fusion Pore and Diffusion

Previous efforts to understand spike shape include the use of an exponentially modified Gaussian (6), as well as a model incorporating matrix dissociation and fusion pore expansion (8). Here we first explore a model that combines diffusion and flux through a static fusion pore. To appreciate the time scales for diffusion it is instructive to consider the expression *χ* = (2*Dt*)^1/2^ for the root-mean-square displacement in one dimension in terms of time, *t*, and the diffusion constant, *D* (6 × 10^−6^ cm^2^ s^−1^ for norepinephrine (29)). For *t* = 0.1, 1, and 10 msec, *χ* = 0.35, 1.1, and 3.5 µm, respectively. Gently touching the electrode to the cell surface should produce an approximately flat contact with a separation of a few tenths of a micrometer. The rise times for amperometric spikes fall within the range of times for diffusion over such distances (Fig. 2B), but the fast decay times are somewhat longer (Fig. 2C; mean 1.84 msec implies *χ* = 2.8 µm). Thus, even the fast component of decay is too slow to be diffusion limited. The following analysis uses a more quantitative treatment of diffusion to model spike shape.

For diffusion to the electrode surface, the site of release is taken as a point source on the surface of a reflecting planar boundary in which *N*_*0*_ molecules have an initial delta function distribution. The electrode is taken as an absorbing planar boundary parallel to the cell surface at a distance of *x*. The solution of the diffusion equation for these conditions gives the rate of absorption by the electrode, which is the amperometric current versus time (2, 30).

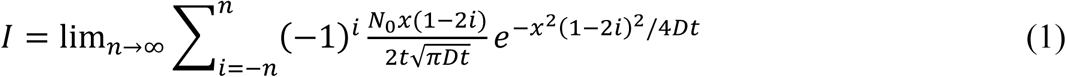

This series converges rapidly; increasing *n* above 5 produces no discernable changes. This expression can produce a range of shapes by varying *x*, but it cannot account for the range of spike shapes with *x* < 1 µm (30). This supports the view that release is not instantaneous and that other processes slow the loss of catecholamine from a vesicle.

To incorporate fusion pore flux we start with a point introduced by Almers et al. (31). If flux through a fusion pore is rate limiting, gradients inside the vesicle are small. Because the electrode is an absorbing surface, the external concentration remains low. The flux through the fusion pore is then proportional to the concentration within the vesicle. The rate of loss of molecules from the vesicle is

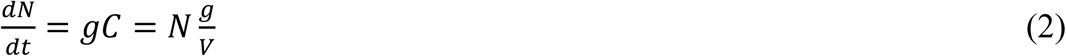

*N* is the number of molecules in a vesicle as a function of time, *V* is the vesicle volume, and *C* is the concentration in the vesicle (the conversion of units is irrelevant at this point but will be addressed as needed). *g* represents the coefficient of proportionality between vesicle concentration and flux, and can be viewed as the fusion pore permeability. *C* and *N* decrease with time after the pore opens, and Eq. 2 gives

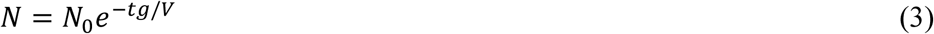

Thus, pore limited flux predicts an exponential decay with a time constant

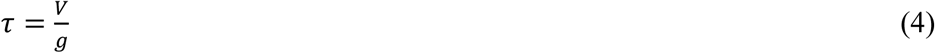

Note that this result depends on a static fusion pore that does not change as the vesicle empties out. This assumption is integral to the analysis of this section and will be relaxed in the ensuing section.

The effects of diffusion and pore flux can be combined by taking the convolution of Eq. 1 and 3.

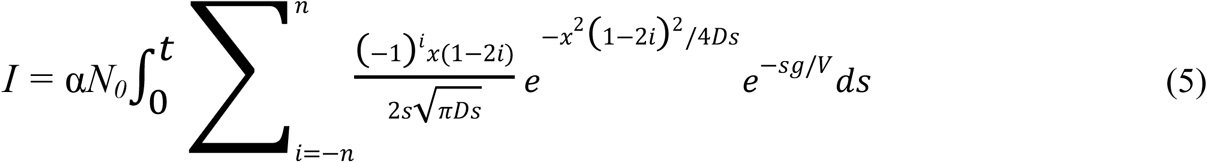

α is 2xFaraday’s constant to convert molecules/s to current. This expression bears some resemblance to the exponentially modified Gaussian function used previously (7), but Eq. 5 treats diffusion explicitly and incorporates a dependence on *x*, the distance to the electrode surface.

Eq. 5 can be used to simulate spikes and explore how parameters influence their shape. Fig. 3A illustrates the impact of varying *N*_*0*_ within the range of observed values (Fig. 2D). Fig. 3B illustrates the impact of varying *x* over the range expected for gentle contact between the electrode and cell. The parameter *g* was fixed at 25 msec^−1^, a value that generates realistic spikes within the ranges used for *N*_*0*_ and *x*. Increasing *x* broadens spikes by slowing both the rise and decay. Increasing *N*_*0*_ also broadens spikes but the effect is entirely on the decay. Thus, for a static pore the decay primarily reflects the emptying of the vesicle, even when diffusion is taken into account. Eq. 1, which ignores the impact of the fusion pore, produces dramatic changes in spike shape when *x* is varied (30). The smaller changes in spike shape in Fig. 3B illustrate how grading the flux through a fusion pore with Eq. 5 dampens the effect of distance.

**Fig. 3.**
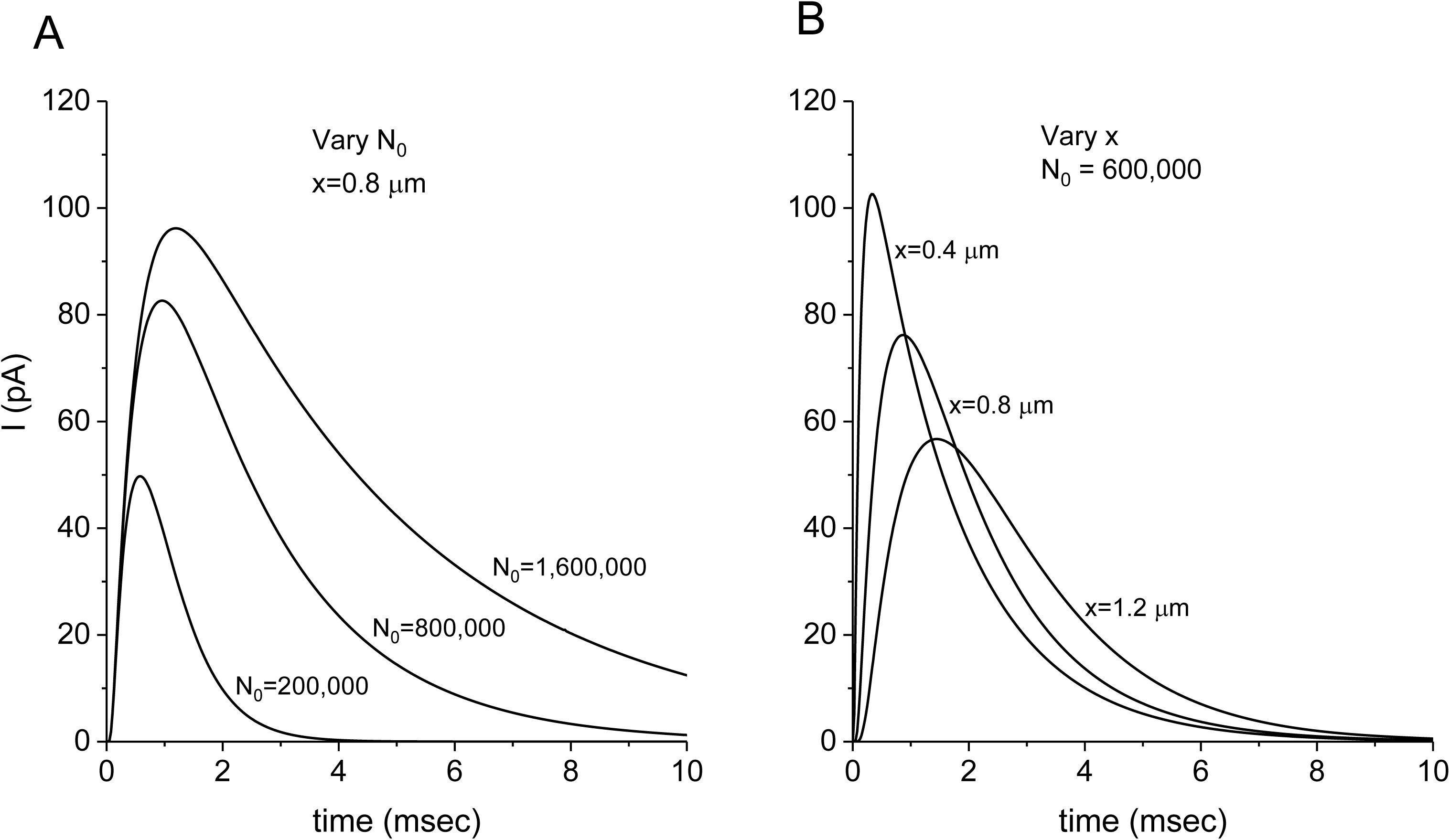
Simulated spikes from the diffusion-static pore model (Eq. 5) for varying *N*_*0*_ with x fixed (**A**) and varying *x* with *N*_*0*_ fixed (**B**). *g/V* = 25 msec^−1^.

The spikes simulated in Fig. 3 by the diffusion-fusion pore model (Eq. 5) resembled the phase of recorded spikes after the pre-spike foot and before the post-spike foot (the “inter-feet” portion). Furthermore, Eq. 5 fitted the rapid components of individual spikes very well. The sum-of-squares error between Eq. 5 and recorded currents was minimized with the Minerr function of Mathcad. To isolate the inter-feet portion, points were taken within 50% of the peak (dashed line in Fig. 4). Spikes were often quite brief, so these fits were performed on data acquired at 10 kHz to provide more points. Eq. 5 has three parameters to vary for the fit, *x, N*_*0*_, and the ratio *g/V*. The onset time was also varied as a fourth parameter because this value is not readily extracted from the data. The start time was generally within a millisecond of the inflection between the foot and spike upstroke. Eq. 5 produced good fits in 140 of 146 events recorded from 3 cells at 10 kHz (Fig. 4, blue curve). The mean parameter values from these fits are presented in the figure legend.

**Fig. 4.**
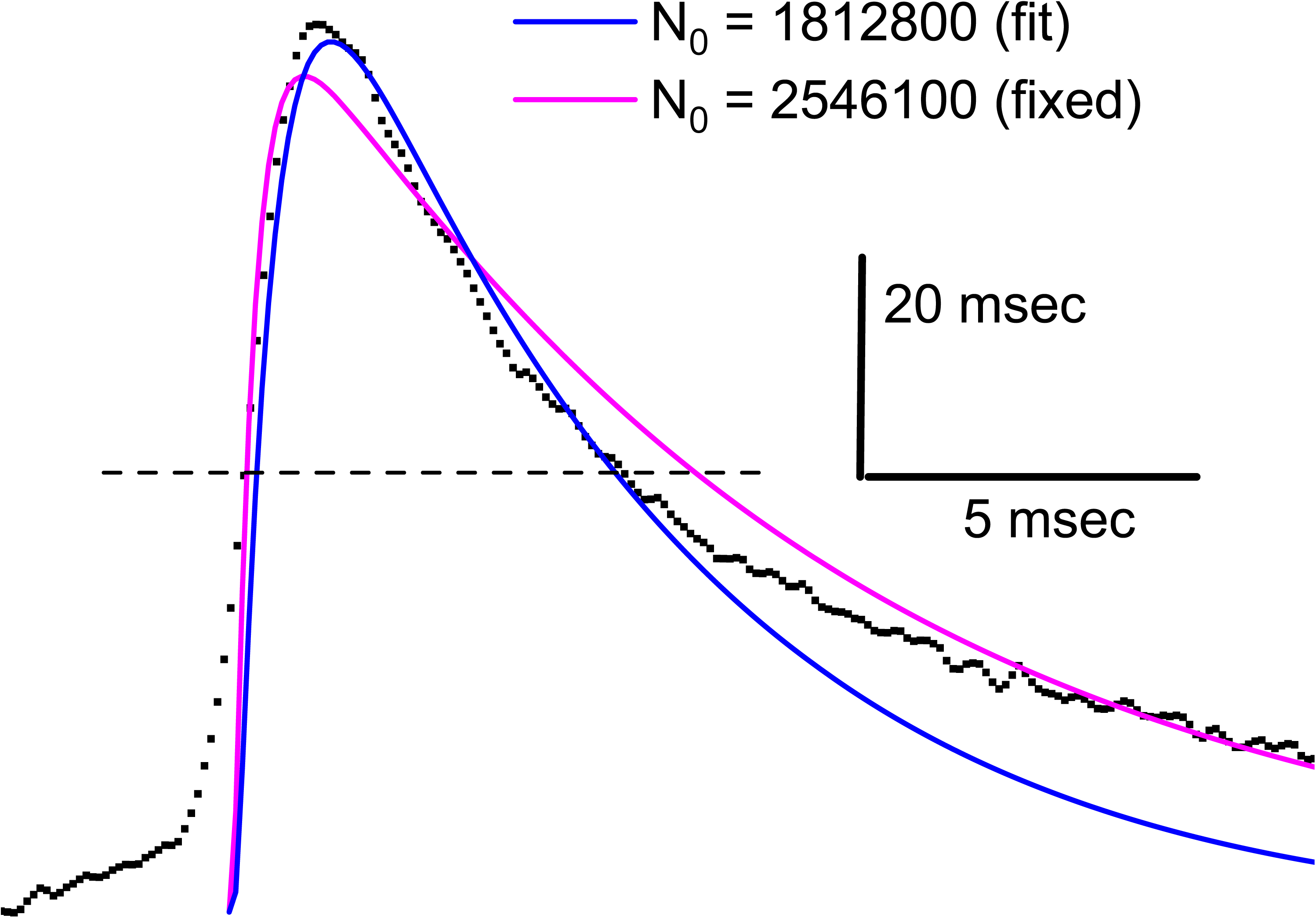
Fits of the diffusion-static pore model (Eq. 5) to the fast part (inter-feet) of a spike. The black points are amperometry data acquired at 10 kHz. The blue trace is the fit with all parameters varied including *N*_*0*_. The magenta trace is the fit with *N*_*0*_ constrained to the value obtained from the area of the event. Fits were conducted on 140 of 146 events from three cells. The 6 discarded events had irregular shapes. Means ± standard deviations. *x*=1.20±0.51 µm; *N*_*0*_ = 1060015 ± 1159924; *g/V* = 136 ± 614 msec^−1^. The mean RMS error of the 140 fits was 0.80 pA.

Although events were fitted well there are telling trends in the parameters obtained. The mean value for *x* of 1.2 µm is larger than the spacing between the electrode and cell. This may reflect an interaction between *x* and other parameters during the fitting process. There was a systematic discrepancy between the value *N*_*0*_ yielded by the fit and the value determined independently from the total area of an event. *N*_*0*_ from all but 4 of the 140 fits was less than the value from the total event area. The mean ratio, *N*_*0-fit*_/*N*_*0-area*_, was 0.68. When *N*_*0*_ was constrained to the value from the area, the fit was very poor (Fig. 4, magenta curve). This represents a meaningful difference between the model and data. A particularly appealing explanation for this result is that 68% of the molecules in a vesicle are free and readily released while 32% are bound to a matrix. This implies that the matrix-bound fraction is responsible for the slower component of decay of the post-spike foot. In fact, post-spike feet accounted for 34% of the spike area on average. Thus, the number of free molecules determined by fitting Eq. 5 is in good agreement with the number contained within the fast component of the spike.

The diffusion-static fusion pore model was tested more critically against experiments by examining the relations between selected indices of spike shape. The time constant for decay (Tau decay) of simulated spikes was plotted against peak amplitude for *x* ranging from 0.3 to 1.2 µm (Fig. 5A). This plot shows Tau decay decreasing weakly with peak amplitude over a modest range. By contrast, varying *N*_*0*_ (for *x* = 0.8 µm; a value that recapitulated the observed mean rise time of 0.5 msec.) yields the opposite dependence; Tau decay increased with peak amplitude over a much larger range (Fig. 5C). Varying *N*_*0*_ shows that peak amplitude increases and gradually reaches a plateau (Fig. 5B), while Tau decay increases linearly (Fig. 5D). This linear increase follows from Eq. 4: for a fixed initial concentration (32), the volume, *V*, is proportional to *N*_*0*_. The preservation of this relation in Fig. 5D underscores the point that grading release through a fusion pore diminishes the impact of diffusion.

**Fig. 5.**
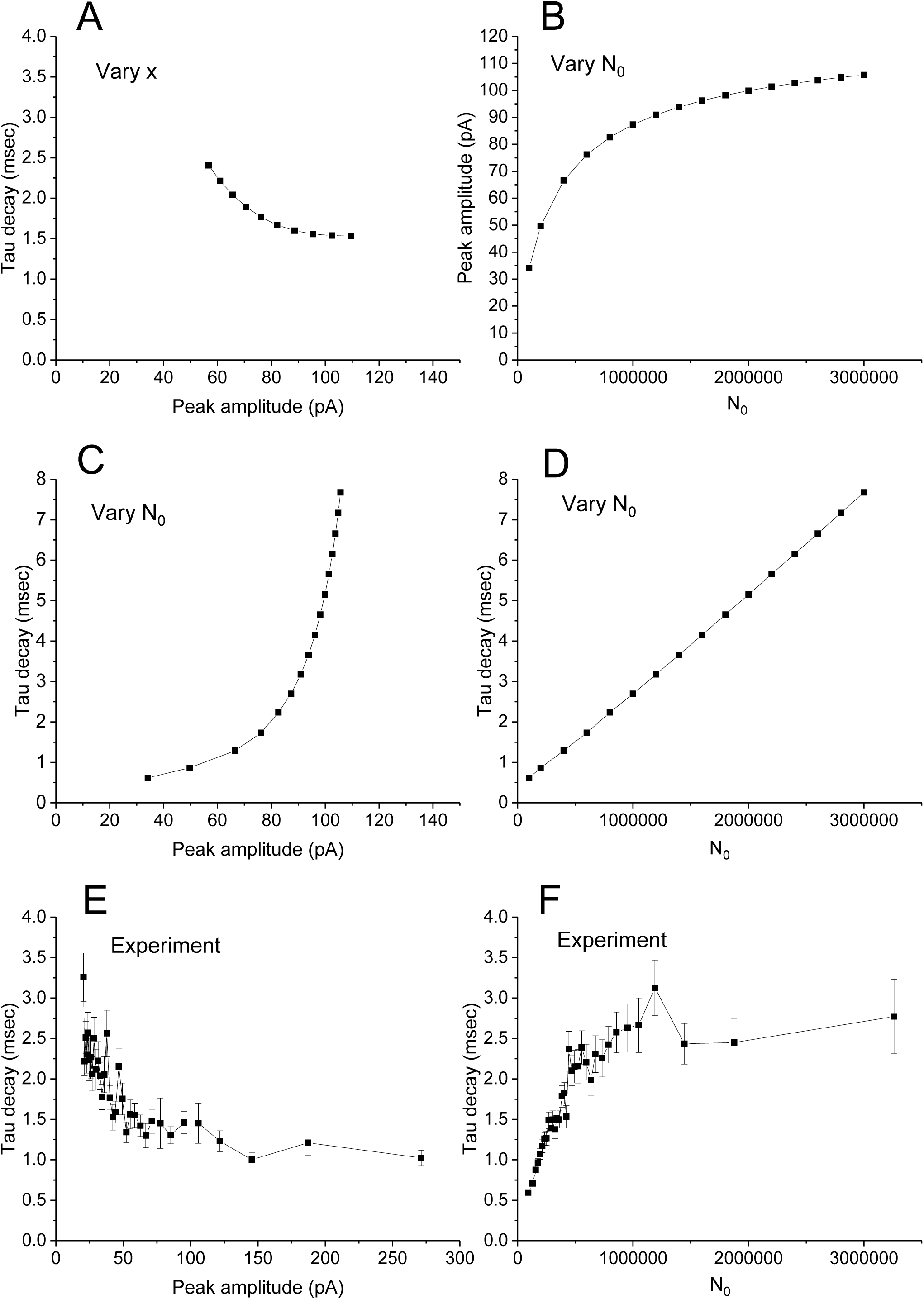
Plots of shape indices from simulations (**A-D**) and experiment (**E** and **F**). **A.** In spikes such as those in Fig. 3 simulated with the diffusion-static pore model, the diffusion distance *x* was varied from 0.3 to 1.2 µm and the decay time was plotted versus peak amplitude. Note that simulated spikes decayed monotonically with a single exponential. **B**. Peak amplitude was plotted versus *N*_*0*_ with x = 0.8 µm. **C**. For the spikes simulated with *N*_*0*_ varied in the range used for **B**, decay time was plotted against *N*_*0*_. **D.** Plot of Tau decay versus *N*_*0*_ with *x* = 0.8 µm. **E.** Plot of Tau decay for the rapid component of versus peak amplitude for 1682 events recorded from 77 cells. Values were binned in groups of 50. Means ± standard errors. **F**. Plot of Tau decay versus *N*_*0*_ determined from the spike area. Numbers and binning as in **E.**

These relations were compared to experiments by constructing plots of peak amplitude, Tau decay (of the fast component, as in Figs. 1B1, 1B2, and 1B3), and *N*_*0*_ from the 1682 events used to construct the distributions in Fig. 2. To view trends clearly, events were grouped into bins of 20 and averaged. A plot of Tau decay of the fast component decreases with peak amplitude (Fig. 5E). The plot based on simulated spikes shows qualitatively similar behavior with varying *x* (Fig. 5A), but the range of both Tau decay and peak amplitude are much wider in the experimental plot than the theoretical plot. To account for these wide ranges, variations in *N*_*0*_ must be considered, but this results in the opposite relation between Tau decay and amplitude of simulated spikes (Fig. 5C) versus experimental spikes (Fig. 5E). The experimental values of Tau decay increase with increasing *N*_*0*_, and saturate with large *N*_*0*_ (Fig. 5F), while the plot from simulations was linear over the entire range (Fig. 5D). The parameter interdependencies predicted by the model differed qualitatively from those observed experimentally. Thus, despite the good fits illustrated in Fig. 4, flux through a static fusion pore cannot account for spike shape. This point was made previously (6, 7), and the present analysis extends this point to a focus on the fast component of spike decay, with a model that treats diffusion explicitly.

### Dynamic Fusion Pores

The preceding analysis of spike shape was based on the assumption of a static fusion pore. Given the shortcomings of this approach just illustrated, the possibility of a dynamic fusion pore was considered. This approach begins with a rearrangement of Eq. 2.

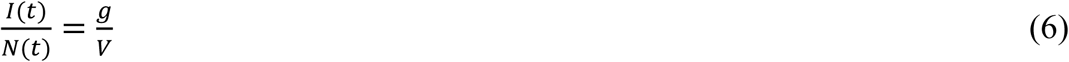

The time dependences of *I* and *N* are now emphasized (again the difference in units of *I* and *N* is irrelevant at this point). Since *N(t)*, the number of remaining molecules, is *N*_*0*_ minus the number lost, and the number lost can be obtained as the integral of *I(t)* from 0 to t, we can express the left-hand side in terms of *I(t)*.

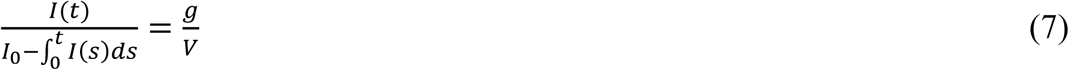

*I*_*0*_ is the total area under the entire event from start to end (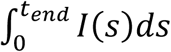; *N*_*0*_ was computed from this quantity). With the numerator and denominator both expressed in terms of current, *g/V* can be calculated versus time for each amperometric event. Eq. 7 thus transforms amperometric current to *g/V*.

Fig. 6 presents examples of *I* and *g/V* (from Eq. 7) for four spikes. These plots exhibit highly characteristic patterns. One important feature is that *g/V* undergoes two major transitions, first with an upstroke at the end of the pre-spike foot, and then a downstroke following the peak in *I*. The initial increase corresponds with a rapid expansion of the initial fusion pore to end the pre-spike foot and start the spike. The rapid decrease after the peak comes as a surprise. Not only does *g/V* decline precipitously, it settles into a stable plateau that lasts tens of milliseconds, for as long as the post-spike foot is visible. The plots become noisy toward the end because the numerator and denominator of Eq. 6 both become small, but *g/V* remains flat for as long as its value can accurately be determined. Thus, the fusion pore remains stable, without closing or expanding, for as long as catecholamine flux can be measured. The transform to *g/V* thus reveals that the post-spike foot (28) corresponds with a stable “post-spike pore”.

**Fig. 6.**
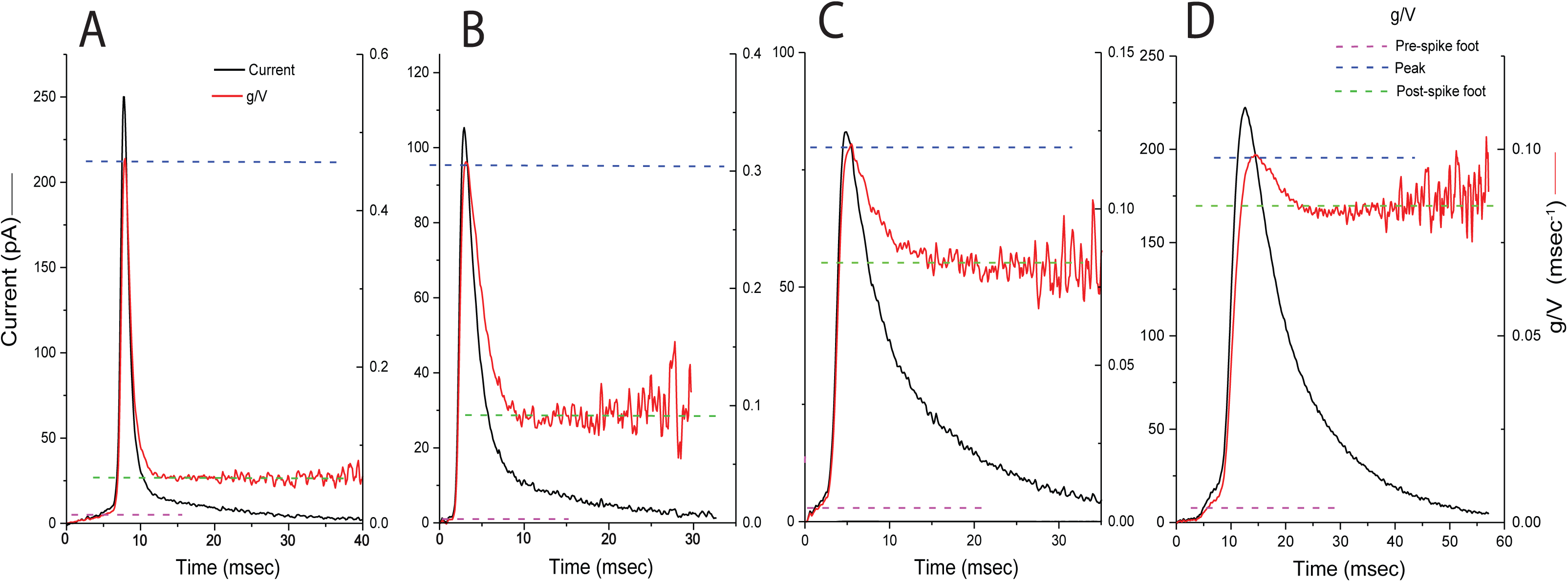
Amperometry spikes (current) were transformed to *g/V* (red) with Eq. 7. In each of the four examples (**A-D**) dashed lines indicate *g/V* of the pre-spike foot (magenta), peak (blue), and post-spike foot (green). The examples illustrate the characteristic initial expansion-contraction cycle of *g/V*, with settling to plateaus of variable amplitude (see text).

The rapid decrease in *g/V* implies that the fusion pore contracts immediately after it expands, and the plateau implies that the fusion pore settles into a stable structure. The three key levels of *g/V*, the pre-spike foot, peak, and post-spike foot, are highlighted in each example in Fig. 6 with dashed horizontal lines as indicated in the legend in the upper right corner. The rapid onset of the spike is likely to reflect both diffusion and pore expansion. The transform to *g/V* does not take diffusion into account, so the actual value of *g/V* at the peak could be larger than the apparent peak in the plots. By contrast, diffusion should play essentially no role in the final plateau of the post-spike pore.

The plots of *g/V* show a large amount of variation in the magnitude of the decline from the peak to the plateau. Fig. 6A declines by ∼90%, Fig. 6B by ∼75% decline, Fig. 6C by ∼30%, and Fig. 6D by ∼10%. Regardless of the extent of decline, a stable post-spike pore was seen in 443 *g/V* plots derived from 449 events recorded from 7 cells. The mean values of *g/V* of the pre-spike foot, peak, and post-spike foot are plotted Fig. 7A to illustrate the changes in *g/V* during the distinct phases of release. From the pre-spike foot to the peak, *g/V* increases ∼15-fold on average, and then declines ∼3.5 fold to the plateau. *g/V* is essentially a time-dependent rate constant that represents the time scale over which a vesicle would lose its content through a static fusion pore of that value. Thus, *g/V* for the pre-spike foot indicates it would take ∼30 msec for loss of 63% of a vesicle’s content. By contrast, the much larger pore at the peak of the plot would only need ∼2 msec for a vesicle to lose the same fraction of content. The peak lasts about this long, so during the peak a vesicle can lose roughly half its content. The intermediate sized pore of the post-spike foot would allow loss of 63% of a vesicle’s content in ∼7 msec.

**Fig. 7.**
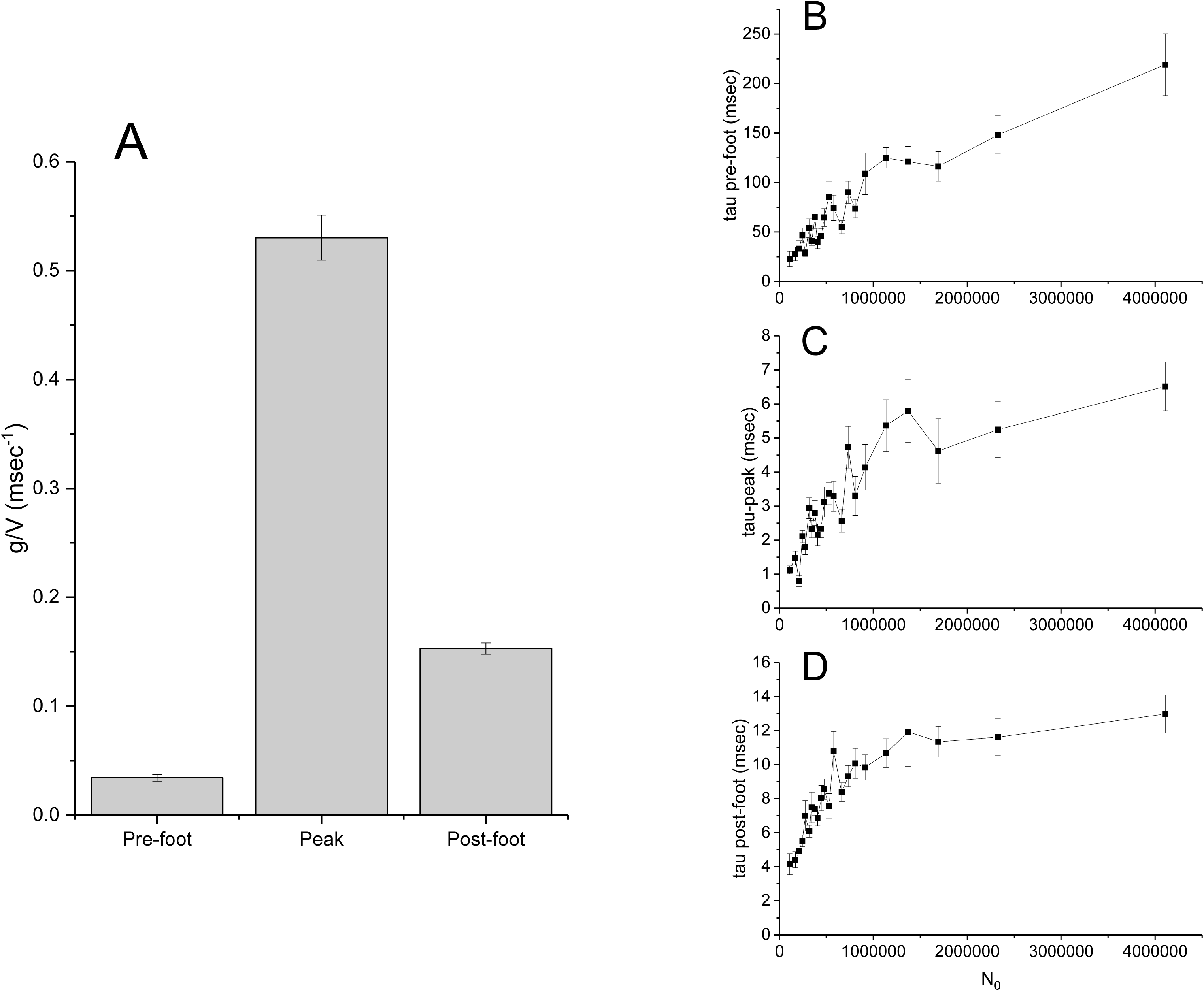
**A**. Bar graph of *g/V* (mean ± standard error) for pre-spike foot, peak, and post-foot. Taking *V/g* as tau, binning in groups of 20 and averaging produced tau plots for the pre-spike foot (**B**), peak (**C**), and post-spike foot (**D**) versus *N*_*0*_. Means ± standard errors.

*N*_*0*_ is proportional to *V* because the catecholamine concentration in a vesicle is independent of vesicle size (32). Thus, we can focus on *g* by examining how *g/V* varies with *N*_*0*_. To relate this analysis to Eq. 4, the reciprocal of *g/V* was taken to give a time constant. For the pre-spike foot the plot of tau versus *N*_*0*_ was fairly linear over the entire range, with an intercept near the origin (Fig. 7B). This suggests a stable fusion pore for the duration of the pre-spike foot, and little variation between vesicles of different sizes. By contrast, for the peak (Fig. 7C) and post-spike foot (Fig. 7D), plots of tau have linear regions for low *N*_*0*_, but the intercepts are above the origin. Furthermore, these tau values reach a plateau for *N*_*0*_ > 1,000,000. This may indicate that pore permeability begins to increase when vesicles are above a critical size. This will be discussed further below.

The transform to *g/V* offers an interesting perspective on the more complex amperometric events that occur infrequently and defy conventional analysis. Most events are spike-like, as illustrated in Figs. 1B1, 1B2, and 1B3, but occasional events with multiple peaks are generally omitted from analysis. Two events with second smaller peaks are displayed in Fig. 8. It is very unlikely that the two peaks in each event are due to coincident fusion of different vesicles, because the overall frequency is too low. The *g/V* traces reveal increases during the second peak, with the fusion pores making a transition to a larger size. The second example (Fig. 8B) shows a further increase after the second peak in current, indicating further expansion. The value of *g/V* becomes noisier and less reliable toward the end of the record due to the problem with quotients of small numbers, but the current trace does show a small departure from monotonic decay starting at about 55 msec. Thus, the fusion pore may actually be expanding further at this point and the *g/V* plot is a better way to detect this.

**Fig. 8.**
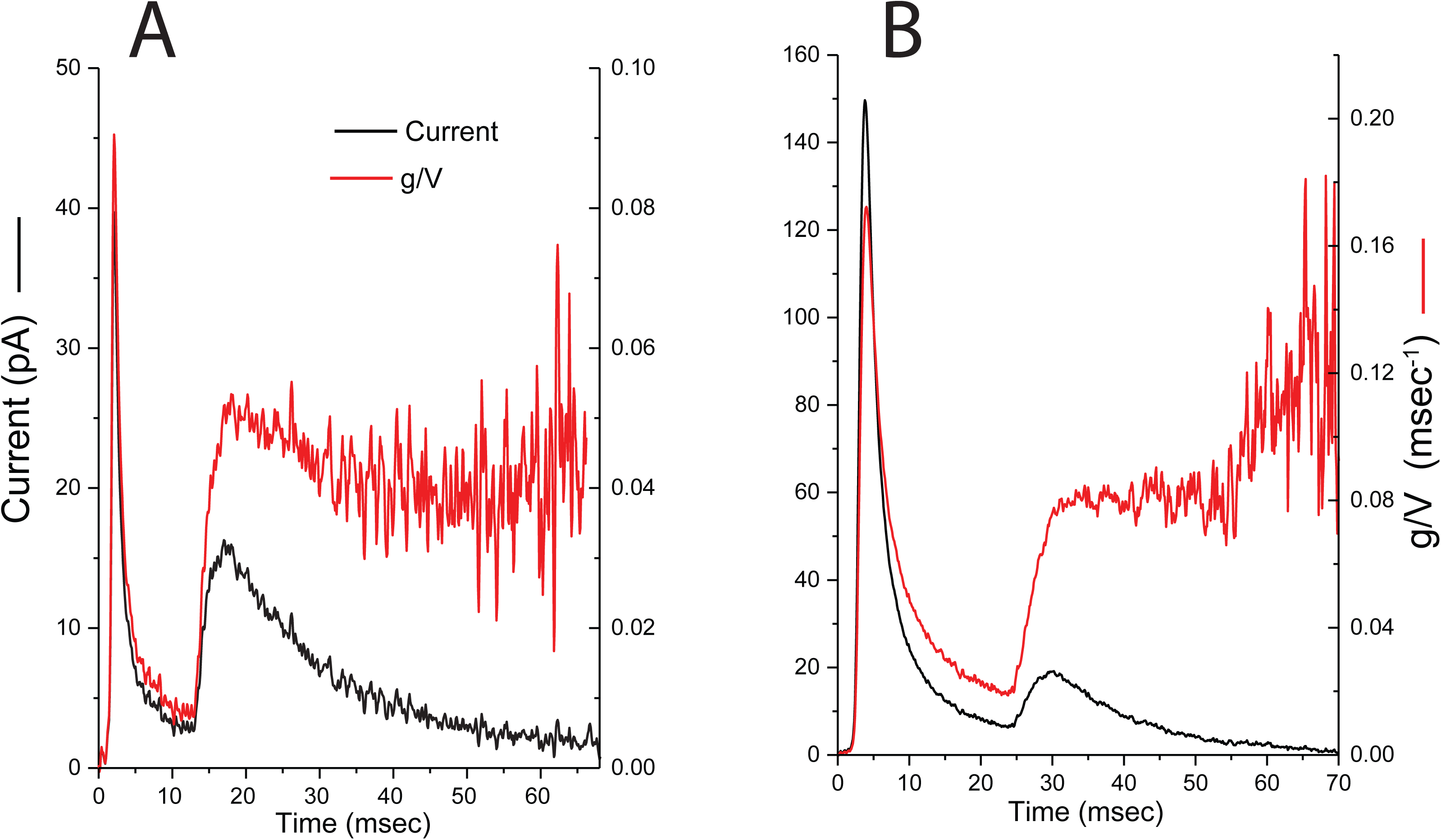
Two examples of current and *g/V* for events with a second peak in amperometric current. The second peak is associated with a stepwise increase in *g/V*.

## DISCUSSION

This study used amperometry data from endocrine cells to investigate the evolution of the fusion pore over time as catecholamine is released from single vesicles. Two different approaches were explored. First, a static pore model was developed which failed to account for important experimental observations. The assumption of a static fusion pore was then relaxed to explore the consequences of dynamic changes in permeability. This yielded a highly characteristic fusion pore permeability time course, beginning with a rapid expansion-contraction cycle, and then settling into a state that is stable for tens of milliseconds. The assumption of a static pore is restrictive, and generates testable predictions. The assumption of a dynamic fusion pore is more robust, and suggests interesting new possibilities. It is significant that the stable plateau was generated by a dynamic model, which makes no assumptions about stability.

### The Failure of Static Fusion Pores

The model of a static fusion pore incorporated diffusion, as represented by Eq. 5. This model simulated realistic spikes (Fig. 3), and fitted the inter-feet portions very well (Fig. 4). However, a more detailed analysis based on plots of indices of spike shape lead to conclusions that challenge the static pore model. The vesicles of endocrine cells vary in size, and *N*_*0*_ is probably the most important source of variation when events are recorded with an electrode gently touching a cell (6). The predicted relation between Decay time and amplitude (Fig. 5C) was opposite to that observed experimentally (Fig. 5E). Furthermore, the Decay time should increase linearly with *N*_*0*_ (Eq. 4 and Fig. 5D), without the saturation seen in the experimental data (Fig. 5F). These disparities argue against the view that flux through a static fusion pore underlies the rapid component of spike decay, and motivated the consideration of dynamic changes in pore permeability.

### Matrix Dissociation

Previous studies suggested that catecholamine dissociation from the vesicular matrix provides a better account of amperometric spike shape than flux through a static fusion pore (6-8). Knocking out the matrix protein chromogranin A reduces the packaging of catecholamine within dense-core vesicles (33, 34). The two components of decay in the present work could be interpreted in terms of the rapid loss of a free pool of catecholamine followed by the slow loss of a matrix-bound pool. However, the failure of the diffusion-static fusion pore model argues against the idea of rapid loss of the free pool through a static fusion pore. Furthermore, matrix dissociation should have a complex time course (7, 8), while the stability of *g/V* during the slow component of catecholamine loss (Fig. 6) supports a role for pore flux. One possibility is that there is a tightly bound pool of catecholamine that dissociates very slowly, but this would be too slow to influence the shapes of spikes over the tens of milliseconds relevant to the present study. A critical analysis of the slow component of decay in terms of quantitative models of matrix dissociation may provide a rigorous test for the role played by the vesicle matrix in catecholamine release kinetics.

### Fusion Pore Dynamics

The static pore model generated results at odds with experiment, and this prompted the consideration of a fusion pore that changes dynamically as release progresses. Some previous studies assumed that fusion pores expand at a constant rate (8, 35). Patch-amperometry recording indicates that the conductance of a fusion pore sometimes increases transiently before a rapid contraction that terminates a kiss-and-run event (36). In the present study, the assumption that an amperometry spike tracks the dynamics of the fusion pore served as the basis for a mathematical transformation of current into the ratio *g/V* (Eq. 7). During most fusion events (Fig. 6) this ratio underwent a transient increase suggesting that fusion pores expand and contract in rapid succession. The contraction terminates with a stable plateau. Furthermore, in the occasional amperometry event with a second peak in current, we find a roughly stepwise increase in *g/V* (Fig. 8). These multi-peak events obviously require more complex mechanisms than static pore flux or matrix dissociation. The *g/V* plots of these events provides a simple interpretation for these events as abrupt increases in pore size.

Diffusion probably alters the shape of the rapid rising phase of the *g/V* plots, but the most likely explanation for the plateau in *g/V* is the formation of a stable fusion pore. This indicates that the post-spike fusion pore is intrinsically stable, and that this structure can resist the expanding forces arising from SNAREpin rotational entropy (37) and membrane tension (38). The possibility that *g* and *V* both vary in parallel to keep the ratio constant cannot be formally ruled out. However, vesicle area changes very little during capacitance flickers (39), and when changes have been observed they are slow (40). Although the flow of lipid label through fusion pores appears to be more rapid, it is still somewhat slower than the times relevant to amperometry recording, and the vesicles appear to maintain their volume during this flux (41, 42). Membrane tension and hydrostatic pressure could drive a burst of content loss as the force balance changes abruptly during pore formation or expansion. This would drive a reduction in vesicle radius, R, at a rate given by the expression *dR/dt* = -*σr*^*3*^*/(6πηR*^*3*^), where *σ* is the membrane tension, *r* is the pore radius, and *η* is the viscosity of the aqueous vesicle lumen (43). With *η* = 1 cP, *r* = 1 nm, and a typical value for σ of .03 dynes/cm (38), a 100 nm vesicle will decrease at a rate of 0.16 nm/msec due to flow through a 1 nm pore. This is far too slow to have an impact within the time scale of these experiments.

### Properties of the Post-Spike Pore

Changes in the shapes of lipidic fusion pores have been predicted previously based on the theory of lipid bilayer elasticity. The earliest lipidic pore represents a state reached by the lowest energy trajectory during pore formation, but this is not the same as the global minimum in the energy landscape (44). Fig. 9 illustrates some possible ramifications of these points. The transition from Fig. 9A to 9B will form the initial proteinaceious pore. This pore transitions to lipid, and Fig. 9C represents the shape most easily reached from the proteinaceous pore. This shape does not represent a global energy minimum, and subsequent changes in shape can lower the energy. A fusion pore can decrease its meridional curvature by becoming bowed (45) or teardrop-like (46), and thus reduce its elastic energy. This will lengthen the fusion pore (Fig. 9D) and thus reduce its permeability. This longer pore with lower energy cannot be formed initially because in order to fuse the membranes must be close together (Fig. 9A). The cylinder can only lengthen after the transition that leads to an initial lipidic pore. The transitions illustrated in Fig. 9 illustrate how the decrease in *g/V* immediately after the peak (Fig. 6) realizes an earlier theoretical prediction (45, 46).

**Fig. 9.**
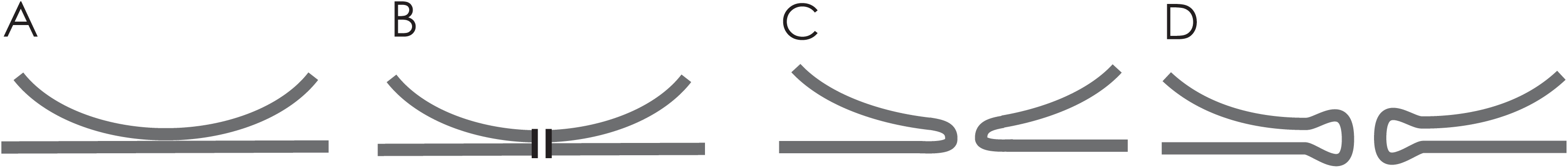
Pore states illustrating changes in elastic membrane bending energies. **A**. A vesicle contacts the plasma membrane to initiate fusion. **B**. The initial pore of the pre-spike foot is formed by proteins and does not alter membrane curvature. **C.** A lipidic pore forms with highly curved membranes and close proximity between the fusing membranes. Creating this state produces a sharp peak in pore permeability. **D.** Membrane elasticity theory has shown that a fusion pore can reduce its membrane bending energy by increasing its length through a reduction in meridional curvature (see text). This shape corresponds with the stable permeability of the post-spike pore

The amperometry data cannot be used to estimate how long the *g/V* plateau lasts because once the catecholamine has left the vesicle there is no signal to report the status of the pore. Single-vesicle capacitance measurements from chromaffin cells indicate that capacitance flickers have durations of hundreds of milliseconds to seconds (26, 47), and these events may represent the continued opening of the post-spike pore. SNARE transmembrane domain mutagenesis also alters the fusion pore conductance during capacitance flickers as well as the amplitudes of pre-spike feet (26, 48), suggesting a role for proteins not only in the small pre-spike pore, but also the larger post-spike pore. The presence of protein in both small and large pores dictates a need for structural flexibility that could be accommodated by a hybrid pore incorporating both protein and lipid (5, 49).

The time constant for the rapid decay of the current exhibited an anomalous dependence on *N*_*0*_ (Fig. 5F), which resembled that of the time constants (expressed as *V/g*) for both the peak and post-spike pore (Figs. 7C and 7D). These time constants all increased with *N*_*0*_, saturated, and had non-zero intercepts. The saturation may indicate that post-spike pores are larger for larger vesicles, or that these pores exhibit less lengthening (Fig. 9). It is also possible that larger vesicles possess a mechanism to actively expel catecholamine. The *N*_*0*_ dependence of *V/g* of pre-spike feet was more linear (Fig. 7B), as expected from Eq. 4, and in accord with the hypothesis of a common pore structure for vesicles of all sizes.

### Revisiting Pre-spike Feet and SNARE Complex Number

The post-spike foot had a *g/V* value 4.5-fold greater than the pre-spike foot (Fig. 7A). Assuming pore permeability scales with area suggests the diameters of the two pores differ by a factor of roughly two. This suggests a significant revision of prior estimates of the number of SNARE protein transmembrane domains that form the initial fusion pore. These studies used the conductance of the fusion pore and the dimensions of an α-helix to estimate that 5-10 transmembrane domains can form a barrel with the appropriate dimensions (48, 50). However, the pore conductance used in these calculations was derived from measurements of capacitance flickers that lasted hundreds of milliseconds. Pre-spike feet, which SNARE mutagenesis experiments suggest reflect flux through pores formed by transmembrane domains (26, 48, 51), last only a few milliseconds. With the factor of two in diameter between the post-spike and pre-spike pores derived from the respective *g/V* values, we can halve the previous transmembrane domain estimate of 5-10 to obtain 2.5-5 SNARE complexes in an initial fusion pore. This brings the number closer to the values estimated for synapses (52) and endocrine cells (53). However, the calculation of pore dimensions based on conductance assumed a fusion pore length of ∼10 nm (two lipid bilayers). The bowing illustrated in Fig. 9D would require a longer pore, with a proportional increase in area. This would raise the estimate of SNARE complex number in the initial pore.

### Biological Functions of the Post-Spike Pore

The post-spike pore may be small enough to retain peptides within vesicles after the onset of fusion (20-22, 54), thus representing an amperometric counterpart to the cavicapture configuration invoked to explain large molecule retention. As noted above, the saturation in the plots displayed in Figs. 5F, 7C, and 7D may indicate that larger vesicles have larger post-spike pores. This leads to the testable prediction that small vesicles should have greater retention of large molecules.

The post-spike pore identifies a potential locus for proteins to influence secretion. Many proteins influence spike shape (9-19). These results have generally been interpreted as an effect on fusion pore expansion, but these studies lacked a framework for relating spike shape to fusion pores. The present analysis provides this framework. Thus, rather than altering pore expansion, the action of dynamin on amperometric spikes (15-17, 19) can be viewed as a change in the size of the post-spike pore or in the speed of the constriction during its formation. This could be related to the collar-forming activity of dynamin around lipidic pores (55). Proteins that alter the post-spike pore will influence the selectivity and extent of content loss. By tracking fusion pores over the entire course of a fusion event, *g/V* derived from amperometric spikes will enhance our understanding of how proteins can control such processes.

## AUTHOR CONTRIBUTIONS

MBJ developed the models, analyzed data, and wrote the paper; YTH performed experiments, analyzed data, and edited the paper; CWC performed experiments and edited the paper.

## ACKNOWLEDGMENT

Supported by NIH grant NS044057.

